# Increased Interactivity and Improvements to the *GigaScience* Database, GigaDB

**DOI:** 10.1101/465666

**Authors:** Si Zhe Xiao, Chris J Armit, Scott C Edmunds, Laurie Goodman, Peter Li, Mary Ann Tuli, Christopher Ian Hunter

## Abstract

With a large increase in the volume and type of data archived in GigaDB since its launch in 2011, we have studied the metrics and user patterns to assess the important aspects needed to best suit current and future use. This has led to new front-end developments and enhanced interactivity and functionality that greatly improves user experience.

In this article, we present an overview of the current practices including the Biocurational role of the GigaDB staff, the broad usage metrics of GigaDB datasets, and an update on how the GigaDB platform has been overhauled and enhanced to improve the stability and functionality of the codebase. Finally, we report on future directions for the GigaDB resource.

## 1. Introduction

There is growing awareness of the societal and technical challenges surrounding reproducibility and rigor in experimental science. The inability of one scientist to reproduce another’s work may be largely attributed to actual artefacts that support the research findings (the data and methods) are largely inaccessible. *GigaScience* (http://GigaScienceJournal.com) is a journal trying to address this by focusing on reproducibility and re-usability when assessing manuscripts rather than subjective impact. Further, we provide citation credit for data to reward and promote the release of large-scale research data for downstream re-use. With data release and credit as a major aim, it is essential that the journal has a data publishing infrastructure that ensures the ability to archive and retrieve the myriad and massive data volumes associated with these ‘Big Data’ publications. *GigaScience* has developed an associated database, *GigaScience* DataBase (GigaDB) (1) to accomplish this and to eliminate the largest excuse for not sharing data: their being too large or that the data types too difficult to make available to the community. GigaDB provides the infrastructure that allows a ‘Big Data’ journal to archive massive multidimensional datasets that are associated with life science and biomedicine studies.

The integration of GigaDB as part of the submission, reviewing and the publishing process, ensures transparency and reproducibility of the article. Importantly, GigaDB is more than just a file server for supplemental files: the datasets are curated by GigaDB staff to ensure the data and metadata are properly curated and organized, and that any links are correct. This assures everything is complete and sufficient for purpose. As a journal with a focus on reproducibility and re-useability of data, peer reviewers have access to all of the supporting data, and the curation process is closely tied to the manuscript review process to ensure timely publication in conjunction with the manuscript, something currently missing from most journal review processes. Trained curators assess the manuscript and data provided by the authors and advise on missing or unclear data and also provide advice on more suitable data formats and recognised standards, such as Genomic Standards Consortium (GSC) (2). Where possible and appropriate, authors are encouraged, and sometimes mandated, to submit data to the relevant public repositories hosted by centres of excellence in the relevant fields, e.g. the European Nucleotide Archive (ENA) (https://www.ebi.ac.uk/ena/) and the Sequence Read Archive (SRA) (https://www.ncbi.nlm.nih.gov/sra).

Datasets hosted in GigaDB are defined as a group of files (e.g., sequencing data, analyses, imaging files, software programs, scripts) and metadata (e.g., sample attributes, protocols, file attributes) that are related to, and support a specific article or study. Through our membership of DataCite (https://www.datacite.org/), each dataset in GigaDB is assigned a digital object identifier (DOI) that should be used as a standard citation for future use in other articles by researchers who use these data. The dataset DOI ensures that the title, creators/authors and additional minimal information are recorded and made permanently available. In addition to fulfilling the requirements of DOI publication, we also aim to maintain the usefulness of the data by allowing minor revisions to be made by tracking all changes in a history log on the dataset page. If major changes are required, a new DOI can be issued and users redirected to the new version with a notice offering to display the original version, which will always be maintained.

GigaDB is managed using the FAIR Principles for scientific data management and stewardship, which state that research data should be Findable, Accessible, Interoperable and Reusable (3). GigaDB datasets are also citable, transforming FAIR into FORCE (FAIR, Open, Research-Object based, Citable Ecosystem (4).

## 2. Datasets, Data Types, and Data Curation

GigaDB hosts datasets from across the life and biomedical sciences and there are currently 20 different dataset types (**Figure 1**). In liaison with the authors of the manuscript GigaDB staff procure all available files and metadata and assist the authors throughout the submission process including assigning dataset types to each dataset. Importantly, multiple types can be ascribed to a single dataset, such that a study that includes both a draft genome assembly (DNA) and aligned transcriptome reads (RNA) would be tagged as both ‘Genomic’ and ‘Transcriptomic’. The GigaDB website utilises these tags and allows users of the web interface to filter datasets based on these categories.

**Figure 1.**
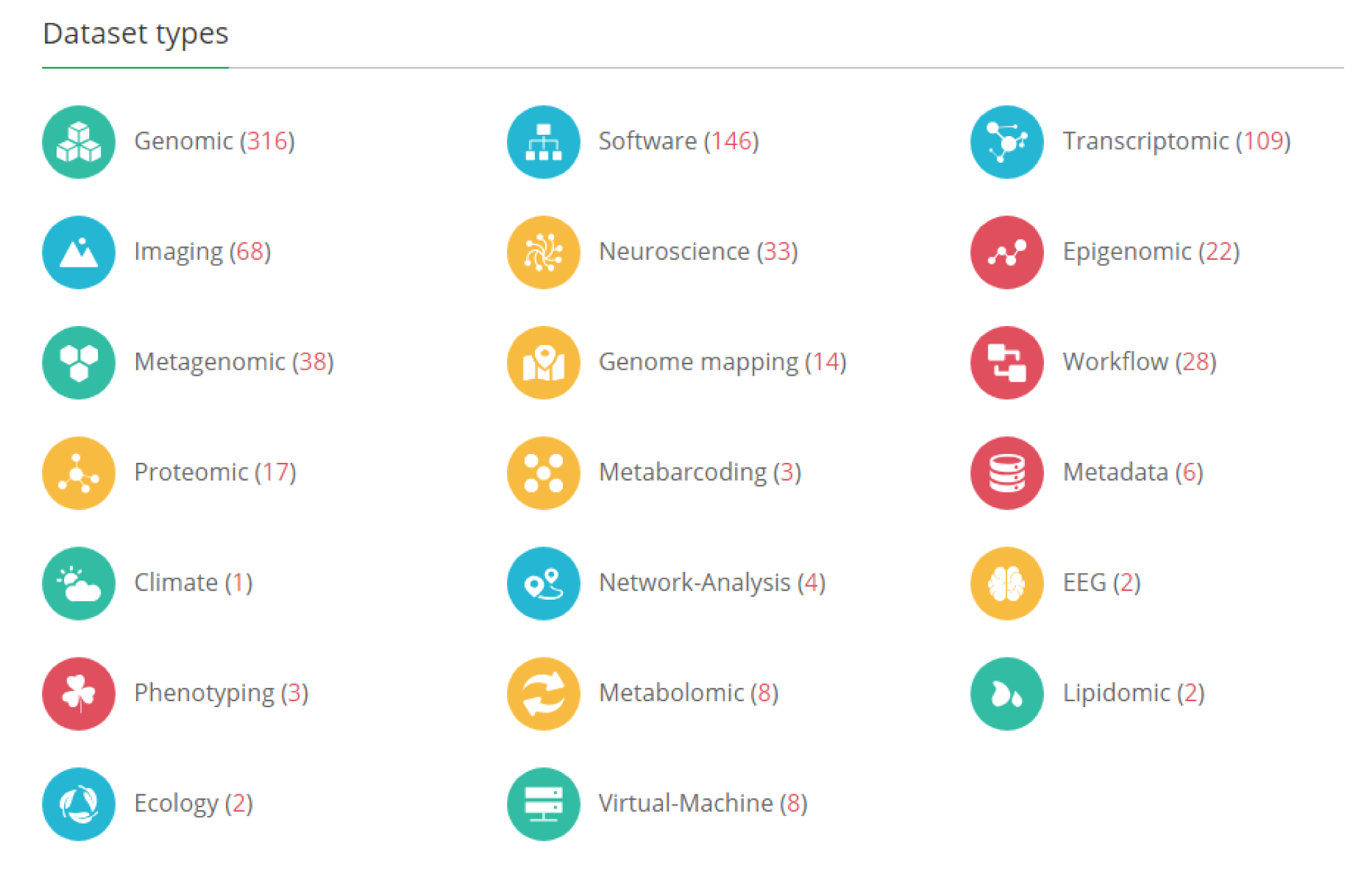
Data Types in GigaDB. The GigaDB home page lists the dataset types that are currently archived in GigaDB. The values in parentheses indicate the number of datasets per dataset type. The end user can click on these links to retrieve these datasets as a list.

As discussed in detail (5) the need to balance the carrot and stick approach to publishing large datasets is still apparent, and many authors are not yet experienced in this matter. GigaDB staff are experienced professional curators, who can provide personalised assistance and guidance to authors where required, and initially authors are provided with guidelines that can be used as a checklist to ensure that all the necessary files associated with the manuscript are submitted (**Table 1**). We believe these checklists will prove to be especially helpful for authors who are submitting data to GigaDB for the first time. Furthermore, these checklists can easily be extended to accommodate new dataset types as *GigaScience* continues to expand its breadth of coverage within the life sciences. Indeed, part of the challenge at GigaDB is remaining cutting edge in the ability to identify the appropriate file types that should be submitted with each new dataset type. Finally, by having the data available for the peer review process, the referees are also able to provide input on which data are important and if there are additional data still missing.

**Table 1.** Guidelines tailored to Dataset types. In the GigaDB guidelines we provide a checklist of files that we would expect to be submitted along with a dataset. This example lists the files that we would anticipate for a genomic and/or transcriptomic dataset. For checklists of other dataset types please see our submission guidelines on our website (https://gigadb.org/site/guide).

| **The expected files for Genome datasets may include;** |  |  |
| --- | --- | --- |
| **Item** | **Suggested format** | **Check** |
| Genome assembly | fasta |  |
| Coding gene annotations | GFF |  |
| Coding gene nucleotide sequences | fasta |  |
| Coding gene translated sequences | protein fasta |  |
| Repeats/transposable elements/ncRNAs /other annotations | GFF |  |
| Gene family alignments (multi-fasta) | multi-fasta |  |
| Phylogenetic tree files (newick) | newick |  |
| BUSCO output file(s) (text) | text |  |
| SNP annotations (VCF) | VCF |  |
| Any perl/python scripts created for analysis process | py, pl, etc |  |
| readme.txt including all file names and brief descriptions | text |  |
| **Additionally, these might be included for Transcriptomic datasets;** |  |  |
| De novo transcriptome assembly | fasta |  |
| Aligned reads | bam |  |
| Expression levels | fpkm table |  |

Where appropriate, GigaDB staff ensure that data are submitted to the public repository, and that those data are made publicly available at the time of manuscript publication. This is an exceptionally important part of the data publishing process that is often missed at other journals as it is not embedded as part of the publishing process. The biocuration team include multiple steps in the submission to publication process that remind authors to submit, for example, sequence data to the Sequence Read Archive (SRA) and then to remind them to have these publicly released prior to publication of the *GigaScience* manuscript that references these datasets.

Checking the data types and availability provides an invaluable opportunity to review the sample attributes associated with the biological samples submitted to the data repositories. Submitting authors are encouraged to ensure that the sample attributes they provide are GSC compliant and assisted by GigaDB curators in the use of ontologies and the correct use of the myriad of possible sample attribute terms. The provision of comprehensive sample metadata ensures the best possible reach of data published in GigaDB and helps the end user find and filter relevant datasets. In addition, interconnectivity of data, and the inclusion of links to external repositories that are referred to in the manuscript is encouraged. These external links can include: RRIDs for software and/or databases; Accession IDs to SRA, European Genome-Phenome Archive (EGA), and mass-spectrometry data (Metabolights and PRIDE), and; DOIs to data hosted in external repositories such as FigShare (https://figshare.com/), Dryad (https://datadryad.org/) or Zenodo (https://zenodo.org/); as well as the use of ontology terms instead of free-text.

GigaDB has additionally installed the community annotation layer “Hypothes.is” within the GigaDB codebase. This allows community-based comments/feedback to be annotated on datasets within each dataset page. We feel that there are key advantages to this overlay approach over the traditional “add comments” section at the bottom of a web page as it facilitates adding specific comments and/or tags to the relevant point in the webpage. Hypothes.is also allows registered users to set up groups so that they can have private discussions about a webpage directly on that page, these comments can later be made public if desired.

### Analytics and Metrics on Use

Launched in 2011, and now with over 550 datasets, GigaDB is able to generate some interesting usage analytics. This data complements the previously published usage analysis of individual high-profile datasets such as the Rice 3K data (7) and the genome of the deadly 2011 German *E. coli* outbreak (8).

While we are indexed by the Web of Science Data Citation Index (https://clarivate.com/products/web-of-science/web-science-form/data-citation-index/), Datamed.org (http://datamed.org), and other services that index our DataCite metadata, it is difficult to collect and present accurate data citation statistics. While CrossRef and DataCite event data does collect data citations in an open format, they only collect data citations correctly cited in the reference sections of indexed publications. This misses any misformatted citations and any links to the datasets referred to in the body of the paper, which from our experience has been the majority of usage. While the FORCE11 Data Citation Principles (which we endorsed at its launch) has promoted the correct practice of citing datasets by DOI in the reference section (4) we have been publishing data since before this was published; and despite our efforts in promotion (9) this is still not the norm for other publishers.

We are able to make use of the services provided by Europe PMC (https://europepmc.org/) and Google Scholar (https://scholar.google.com/) to track the number of “citations” (or hits) of GigaDB DOIs in published articles. Google Scholar has a broad reach and (unlike Europe PMC) generally includes articles on the periphery of scientific publishing, including some scientific blog posts, patents and news articles, hence in general the number of citations being returned by Google Scholar is higher than that returned by Europe PMC. In fact every dataset published in GigaDB contains links to the search results from both Europe PMC and Google Scholar, which allows users to get up to date citations from those sources.

By searching for the unique DOI prefix “10.5524/” we have polled Google Scholar and Europe PMC search tools by year, to determine the overall number of appearances of GigaDB datasets in the scientific literature (**Figure 2**). As the trend lines indicate there has been a steady increase in citation of GigaDB datasets over time.

**Figure 2.**
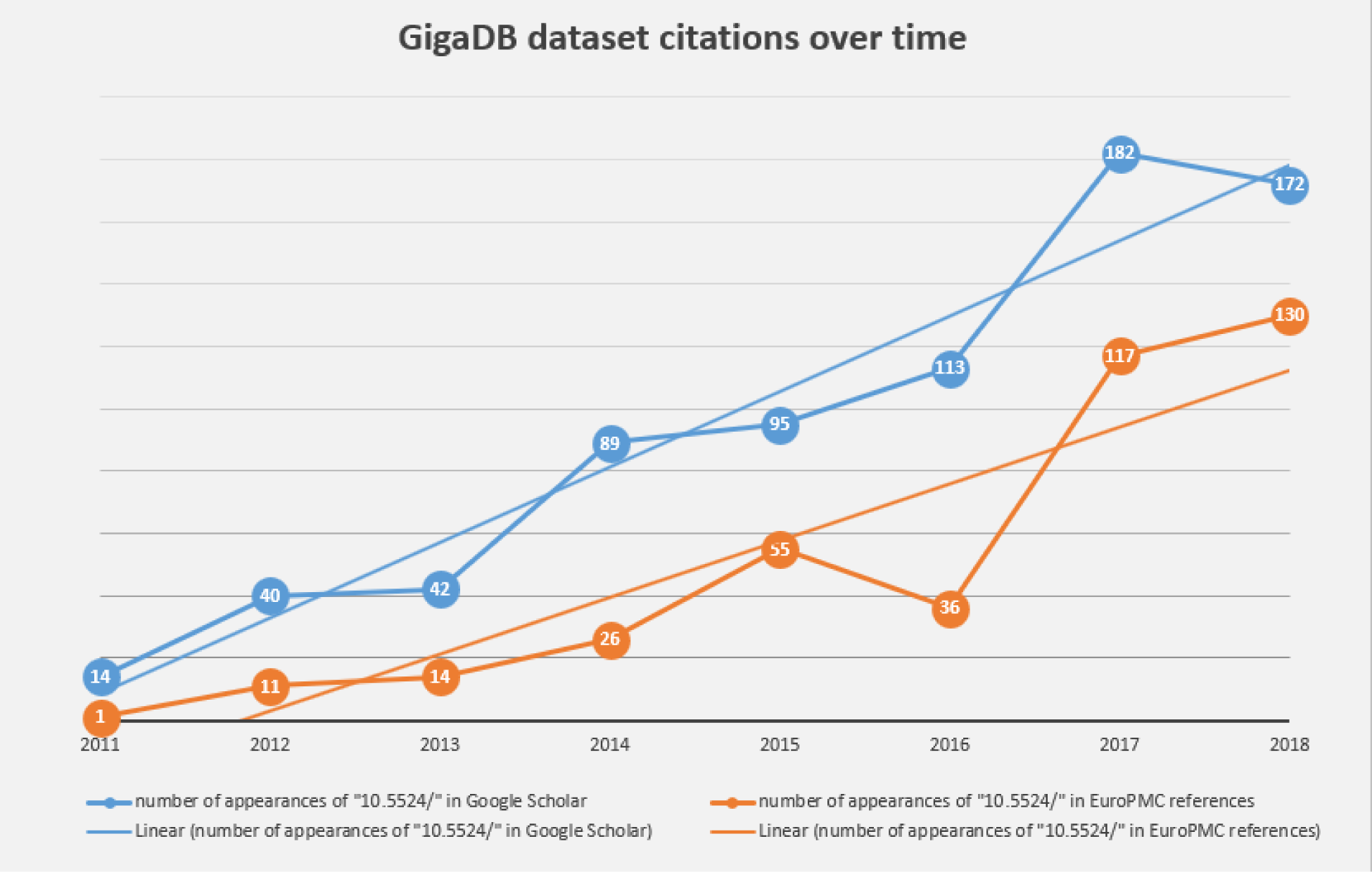
Chart showing steady increase in the number of correctly cited GigaDB datasets found in the scientific literature by Google Scholar (blue) and Europe PMC (orange).

As described above, the FORCE11 Data Citation Principles are not always adhered to, so looking for dataset citations in the literature tells only part of the story. We would miss those instances where authors simply acknowledge a dataset by name within the text, and of course there are likely to be other instances of reuses of datasets that have yet to reach publication. In an attempt to identify more “of the moment” metrics, GigaDB collects other metrics of data usage, including web-browser, data access and file download statistics, as an additional proxy to the official data (re)use metrics.

Google Analytics (https://analytics.google.com/) is a web analytics service offered by Google that tracks and reports website traffic. GigaDB first started using this service in May 2013, and the chart below (**Figure 3**) shows the type of information that can be obtained using this mechanism: in this case, the count of unique visitors per month over the 5 year period 2013-2018.

**Figure 3.**
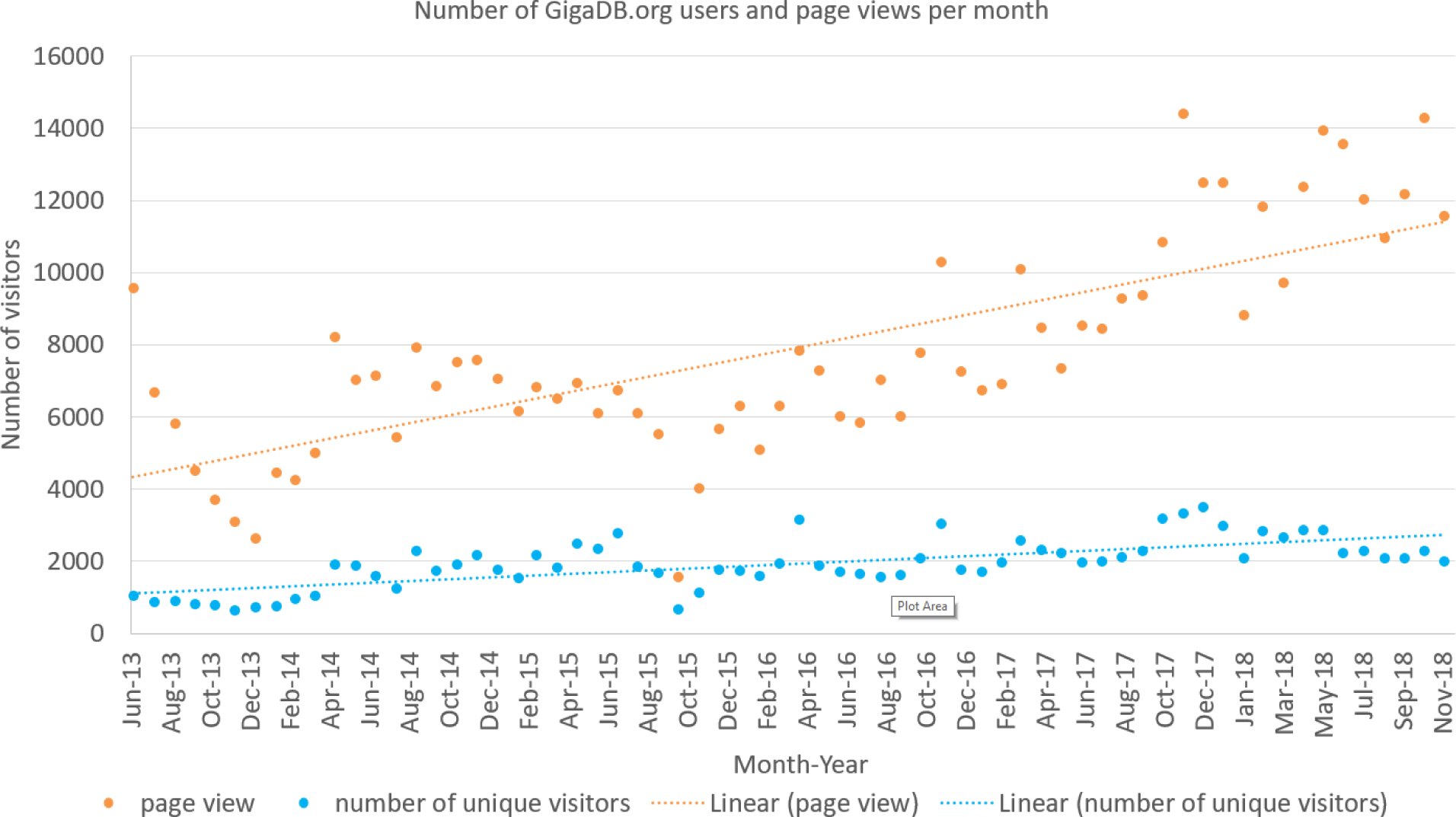
The Google analytics data (June 2013 through to Oct 2018) for the number of unique visitors (blue) to GigaDB.org as well as the total number of page views by all users per month (orange).

Further analysis of Google analytics can be performed on individual datasets to display the number of users browsing any given dataset. Currently this functionality is only available upon request, but we hope to enable this feature in the public view in a future release of the GigaDB code during 2019.

In the period 31st Jan 2017 to 7th May 2018 we analyzed FTP server logs to see how much data was being downloaded. The chart (Figure 4) shows the number of unique IP addresses (used as a proxy for individual users) successfully downloading 1 or more files from any given dataset during that period. On average at least 1 file was downloaded from each dataset by 45 different users during the 15 month period monitored. Some datasets had 10 times that number e.g 100064 (10) was downloaded by 497 unique IPs. The long tail on the chart demonstrates that there are a number of datasets that were downloaded only a handful of times.

**Figure 4.**
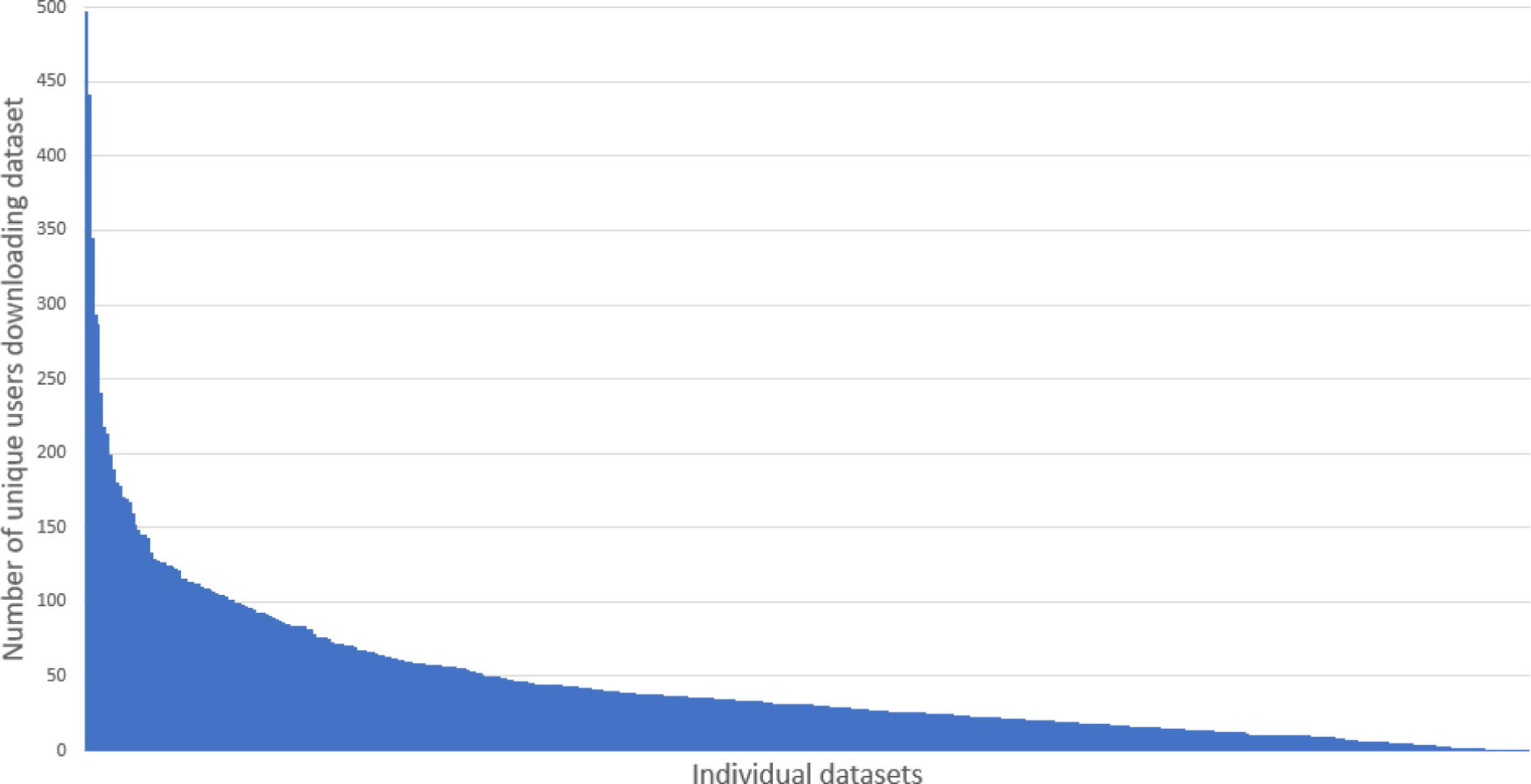
Demonstrates the small number of highly downloaded datasets and the long tail of datasets with fewer downloads.

The FTP log file also records the size of files that are being downloaded, and using those data it is possible to see an increase in data download volume over the period analyzed (**Figure 5**).

**Figure 5.**
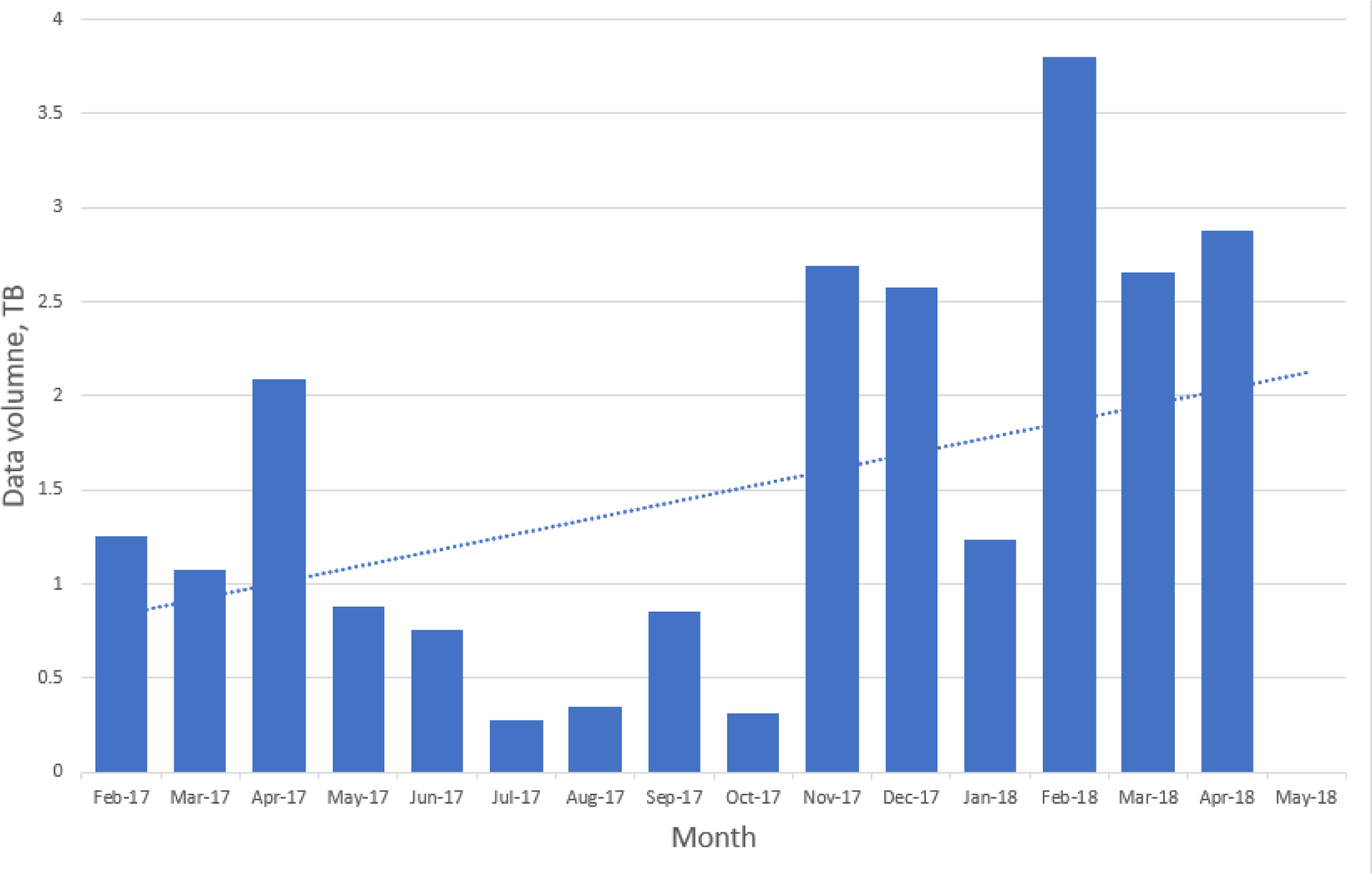
Chart of the overall downloaded data volumes (via FTP) over time

GigaDB have recently been made aware of the “Code of practice for research data usage metrics release 1” (9) that will bring some cohesion to this area, and we fully intend to adhere to the guidelines where possible and participate in the future developments of those guidelines.

## Keeping up appearances: new GigaDB web-based functionalities

Since the last major update in 2014, the GigaDB interface has received an overhaul to provide a more user-friendly dataset page with greater flexibility. On the dataset page, we still maintain the functionality of being able to link to related GigaDB datasets through use of ‘*IsPartOf*’ and ‘*HasPart*’ relationships. However, we now provide multiple tabs as a means of navigating between the GigaDB Sample, Files, Funding, and History information (**Figure 6**). The inclusion of multiple tabs ensures easy navigation and additionally allows novel views of the data. To illustrate with one example, microCT data files can be inspected as a file list as before, but example data files can now also be explored in the context of an interactive 3D viewer (**Figure 7**). Behind-the-scenes, the GigaDB team are generating surface reconstructions for recently submitted microCT datasets and making these publicly available using SketchFab (https://sketchfab.com/GigaDB). The newly launched GigaDB website is able to embed these outside resources within an additional tab on the GigaDB page, and this allows alternative views of the data. Other embeddable media that we currently support are: JBrowse (Genome browser); Code Ocean (actionable software code); and Protocols.io (Methods). A key advantage of the new GigaDB interface is that it allows users this additional functionality within the context of the GigaDB dataset page. Users have the option to pop-out the view into a new window if they so desire.

**Figure 6.**
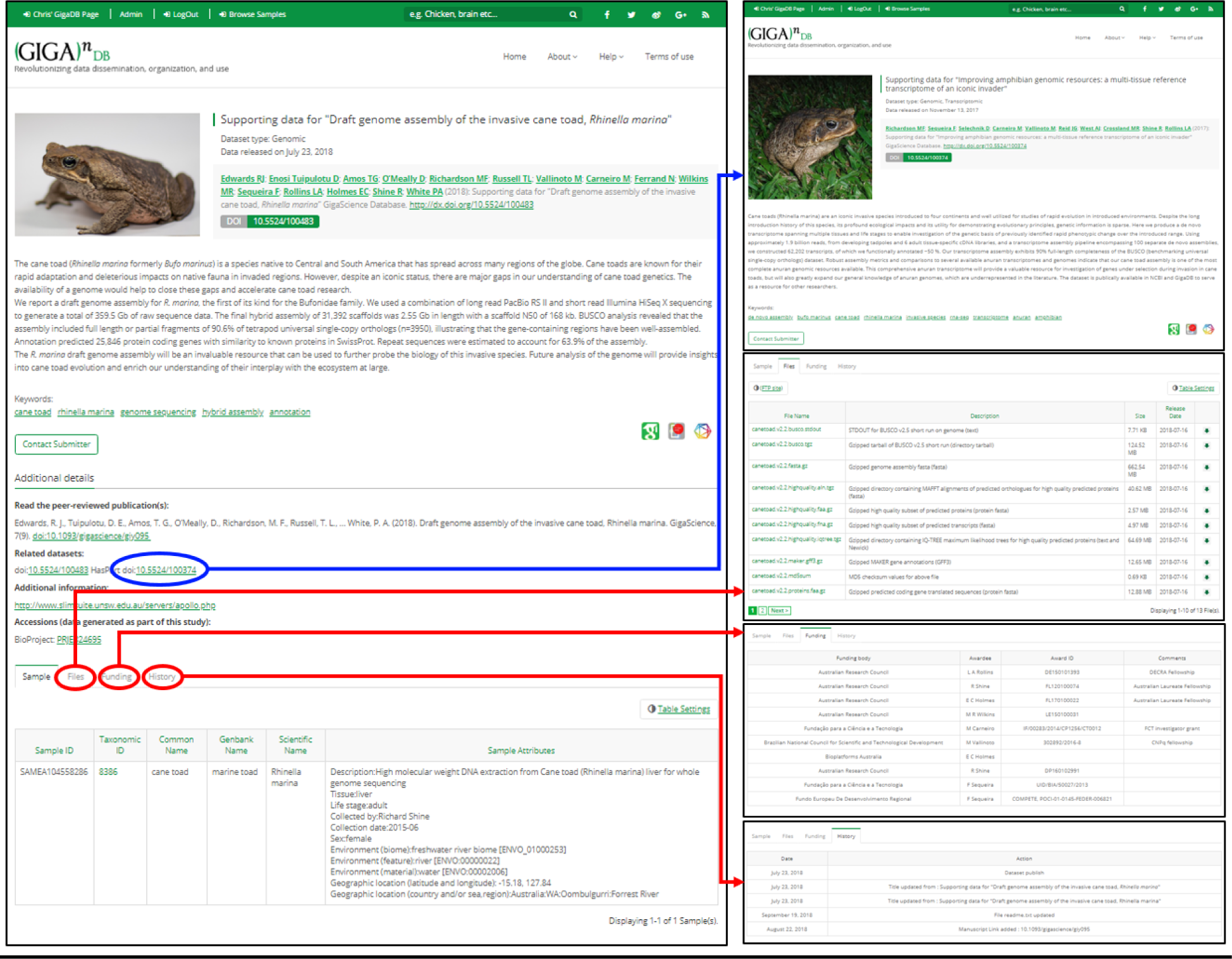
Anatomy of a GigaDB dataset. The dataset page (left panel) includes title, authors, abstract, citation (including DOI), a thumbnail image, and additional links. A key feature is the provision of links to the dataset page of related GigaDB datasets (blue arrow). In addition, a new tab interface allows users to toggle between Samples, Files, Funding, and History (red arrows). The Sample tab is shown by default.

**Figure 7.**
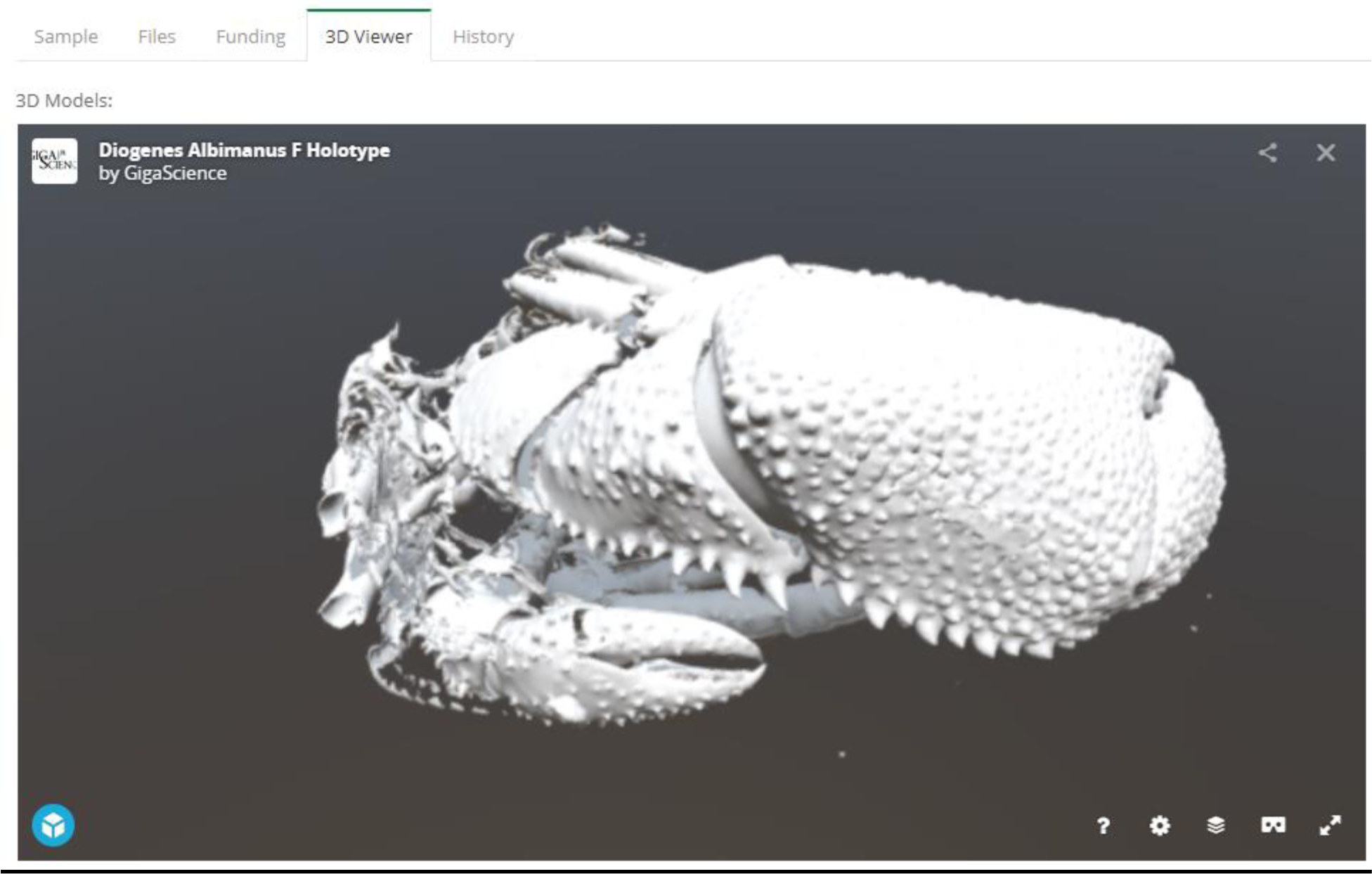
Embeddable media in GigaDB. The GigaDB team are generating surface reconstructions for recently submitted microCT datasets and making these publicly available using SketchFab, example shown from dataset 100364 (11).

We have additionally launched an interactive Map Browser (http://gigadb.org/site/mapbrowse) based on the OpenLayers (https://openlayers.org/) open source JavaScript library for displaying map data that provides a global overview of the datasets that are in GigaDB. The Map Browser uses the latitude / longitude coordinates provided in the sample metadata to generate a point annotation on a world map, with the number of samples assigned to a given location clearly labelled (**Figure 8**). The end user can pan-and-zoom on the Map Browser, and by selecting a point annotation can view a list of samples associated with a specific location and can open these GigaDB datasets in a new tab.

**Figure 8.**
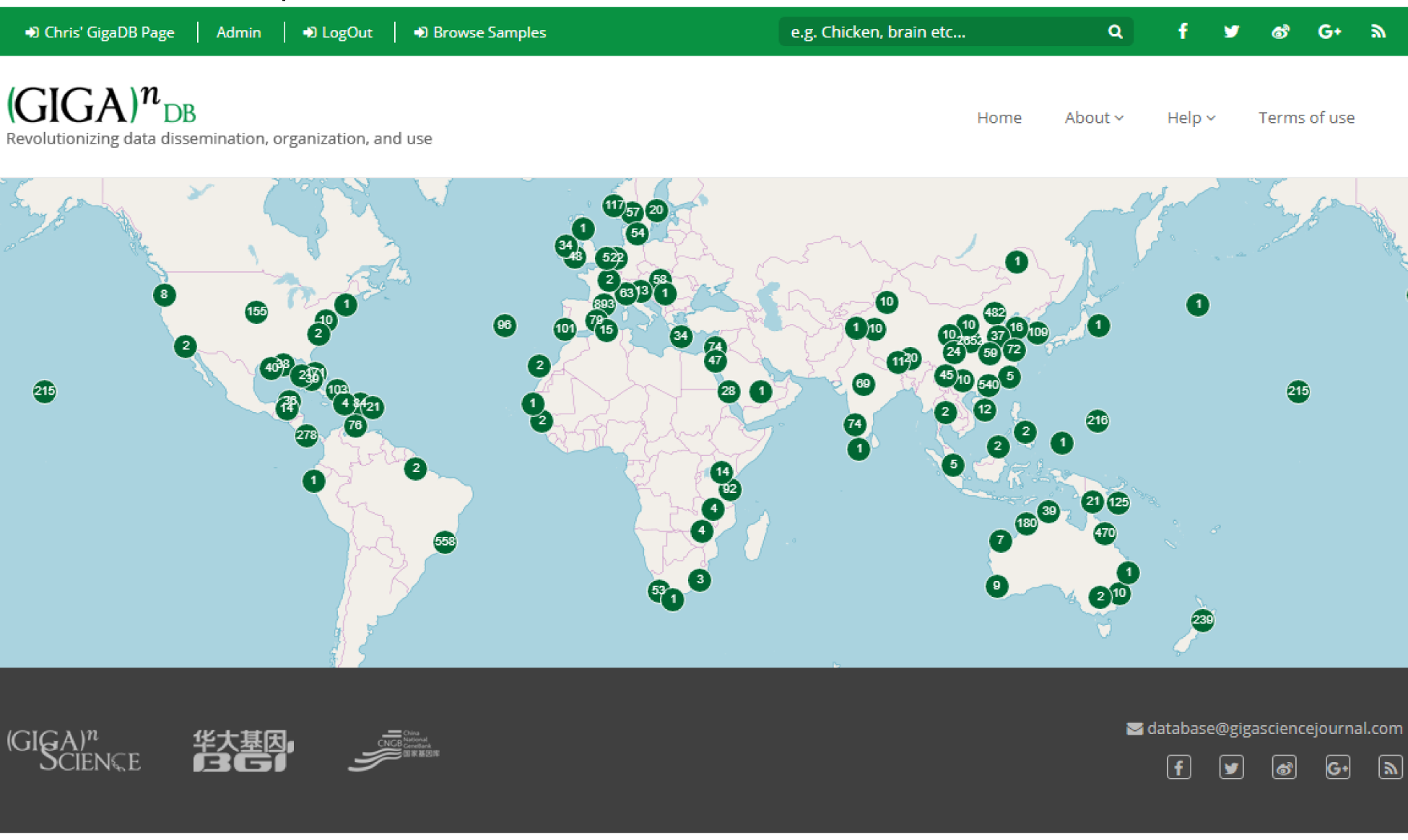
Map Browser in GigaDB. The map browser uses latitude / longitude coordinates to generate point annotations on a world map, with the number of samples assigned to a given location labelled as values. The end user can pan-and-zoom the map, and by selecting a point annotation can view a list of samples associated with a specific location.

### Under the Hood: Improvements to the Code, Testing, and Deployment

The PHP source code for GigaDB is freely-available on GitHub (see availability of supporting source code and requirements). Requests for new functionality and notification of software bugs can also be raised as issues on the GigaDB GitHub repository webpage. GigaDB was initially created with PHP 5 using version 1 of the Object-Oriented, Model-View-Controller Yii framework but recent development work has upgraded it to use PHP 7 and Yii 2.0. The GigaDB source code has also been refactored to reduce its complexity. In addition, a Behavior-Driven Development process is now coupled with Test-Driven Development and unit testing techniques for developing the GigaDB application. We have also adopted Continuous Integration and Continuous Deployment approaches to automate checking of the GigaDB source code so that the application is not broken and that new functionality is made available in a timely manner on the web site, respectively.

## Future directions

### Embeddable media

Whilst we already have in place the Web3D application SketchFab, that allows interactive views of 3D images, we will also provide a section viewer that allows the end user to choose a 2D section through a 3D image and pan-and-zoom on the selected image. The focus here is on developing web-tools that allow researchers to preview large data prior to data download. With minor modification, the section viewer can additionally accommodate time series data, such as time-lapse video microscopy data and camera-trap images, examples of which already exist in GigaDB.

### Submission Tools

To ensure more rapid submission of data associated with *GigaScience* articles, GigaDB is developing a new version of the GigaDB submission wizard. This web-based interface allows authors to submit a new dataset, directly linked to their manuscript submission, online using web-based forms. The aforementioned checklists of data types associated with datasets categories will be integrated into the wizard to facilitate the process. These developments are underway with the intention to release in mid 2019.

## Conclusions

GigaDB have continued to develop its resources further enabling the open sharing of scientific data. The overhaul of the underlying codebase will ensure that developments can continue in a timely manner allowing new and proof-of-concept ideas to be deployed in GigaDB.

The collection of a variety of metrics has allowed monitoring of growth over the recent past and moving forward will provide a solid platform to provide users with vital feedback about the use and citation of their data.

The aesthetic changes to the appearance of the website have enabled the inclusion of widgets which in turn allows for a greater versatility. As always, user feedback is encouraged either by direct interaction with the *GigaScience* and GigaDB staff or via the various online communications.

### Availability of supporting source code and requirements

- Project name: GigaDB: The *GigaScience* Database
- Project home page: https://github.com/gigascience/gigadb-website
- Operating system(s): Platform independent
- Programming language: Python, Java, PHP, postgresql
- Other requirements: PHP 5.6 or higher, Postgresql 9.1 or higher, Nginx 1.4 or higher
- License: GNU GPL v3
- RRID:SCR_004002
- FAIRSharing: doi:10.25504/fairsharing.rcbwsf

## Abbreviations

DOI: Digital Object Identifier
EGA: European Genome-Phenome Archive
ENA: European Nucelotide Archive
FTP: File Transfer Protocol
GigaDB: *GigaScience* Database
GSC: Genomics Standards Consortium
G10K: Genome 10K (10,000 Vertebrate Genome)
IP: Internet Protocol
microCT: Micro computed tomography
SRA: Short Read Archive
PHP: Hypertext Preprocessor scripting language

## Acknowledgements

The authors wish to thank Ernest Lam for the preliminary analysis of the FTP logs for data download usage statistics.

## References

1. Sneddon,T.P., Zhe,X.S., Edmunds,S.C., et al. (2014) GigaDB: promoting data dissemination and reproducibility. Database. DOI:10.1093/database/bau018

2. Field,D., Garrity.G., Gray.T., et al (2008) The minimum information about a genome sequence (MIGS) specification. Nature Biotechnology. 26, 541–547.

3. Sansone,S.A., McQuilton,P., Rocca-Serra,P., et al. (2018) FAIRsharing, a cohesive community approach to the growth in standards, repositories and policies. bioRxiv., DOI:10.1101/245183

4. Martone,M. (2014) Data Citation Synthesis Group: Joint Declaration of Data Citation Principles., DOI:10.25490/a97f-egyk

5. Edmunds, S. C., Li, P., Hunter, C. I., et al. (2016). Experiences in integrated data and research object publishing using GigaDB. International Journal on Digital Libraries, 18(2), 99–111. DOI:10.1007/s00799-016-0174-6

6. The 3000 Rice Genomes Project (2014): The Rice 3000 Genomes Project Data. GigaScience Database. DOI:10.5524/200001

7. Li D; Xi F; Zhao M; Chen W; Cao S; Xu R; Wang G; Wang J; Zhang Z; Li Y; Cui C; Chang C; Cui C; Luo Y; Qin J; Li S; Li J; Peng Y; Pu F; Sun Y; Chen Y; Zong Y; Ma X; Yang X; Cen Z; Song Y; Zhao X; Chen F; Yin X; Rohde H; Liang Y; Li Y; the Escherichia coli O104:H4 TY-2482 isolate genome sequencing consortium (2011): Genomic data from Escherichia coli O104:H4 isolate TY-2482 BGI Shenzhen. http://dx.doi.org/10.5524/100001

8. Edmunds, S. C., Pollard, T. J., Hole, B., & Basford, A. T. (2012). Adventures in data citation: sorghum genome data exemplifies the new gold standard. BMC Research Notes, 5(1), 223. DOI:10.1186/1756-0500-5-223

9. Fenner, M., Lowenberg, D., Jones, M., et al. (2018). Code of practice for research data usage metrics release 1. PeerJ. DOI:10.7287/peerj.preprints.26505v1

10. Li, J., Jia,H., Cai, X., et al. (2014). Supporting data for the paper: “An integrated catalog of reference genes in the human gut microbiome”. GigaScience Database. http://dx.doi.org/10.5524/100064

11. Anton, D. P., Charles, G., & Jannes, L. (2018). Supporting data for “A micro X-ray computed tomography dataset of South African hermit crabs (Crustacea: Decapoda: Anomura: Paguroidea)”. GigaScience Database. http://doi.org/10.5524/100364

